# Expanded methylome and quantitative trait loci detection by long-read profiling of personal DNA

**DOI:** 10.1101/2024.03.17.585420

**Authors:** Cristian Groza, Bing Ge, Warren Cheung, Tomi Pastinen, Guillaume Bourque

## Abstract

Structural variants (SVs) are omnipresent in human DNA, yet their genotype and methylation status is rarely characterized due to previous limitations in genome assembly and detection of modified nucleotides. Because of this, the extent to which these regions act as quantitative-trait loci is also largely unknown.

Here, we generated a pangenome graph summarizing the SVs in 782 de novo assembled genomes obtained from the Genomic Answers for Kids rare disease cohort, that captures 14.6 million CpGs in DNA segments that are absent from the CHM13v2 assembly (SV-CpGs), expanding their number by 43.6%. Next, using 435 methylomes from the same samples, we genotyped a total of 7.99 million SV-CpGs, of which 5.18 million (64.8%) were found to be methylated (SV-5mCpGs) in at least one sample.

To understand the provenance and impact of these novel SV-CpGs, we noted that non-repeat sequences were the leading contributor of SV-CpGs (3.3 × 10^6^), followed by centromeric satellites (1.58 × 10^6^), simple repeats (1.19 × 10^6^), Alus (0.67 × 10^6^), satellites (0.39 × 10^6^), L1s (0.27 × 10^6^), and SVAs (0.19 × 10^6^). Meanwhile, the methylation rate of SV-CpGs was the highest in repeat sequences. Moreover, in contrast to Alus and L1s, centromeric satellites, simple repeats and SVA sequences were overrepresented in SV-5mCpGs compared to reference CpGs. Similarly, we established that non-reference CpGs were more than twice (37% vs. 15%) as likely to be variable, showing intermediate methylation levels in the population.

Lastly, to explore if SVs detected in this pangenome are potentially causal for functional variation in population we measured methylation quantitative trait loci (SV-mQTLs) using CHM13v2 as a backbone. This revealed over 230,464 methylation bins within 100 kbp of a common SV (>5% MAF) showing significant association (at 5% FDR) with methylation variation. Finally, we assessed how many of these SVs-mQTLs were the leading QTL variant compared to SNVs and identified 65,659 methylation bins (28.5%) where the leading variant was an SV.

In conclusion, our results demonstrate that graph genome references providing full SV structures in combination with the associated methylation variation reveal tens-of-thousands of QTLs that are more accurately mapped in personal genomes, underscoring the importance of assembly-based analyses of human traits.

## Introduction

The completion of the first telomere to telomere genome (Nurk Sergey et al. 2022) also enabled the first epigenomic characterization of a complete human genome (Gershman et al. 2022). This milestone epigenome characterized histone modifications and DNA methylation in previously unsolved and structurally polymorphic regions of the human genome, including centromeres, transposable elements and tandem repeats. More generally, the DNA sequences omitted from current reference genomes are likely a source of substantial epigenetic activity. Expanding the non-reference results to a larger number of human genomes and epigenomes can expose population variation with potential new insights on trait variation and disease. The ability to survey epigenomes was recently augmented by long-read technologies that simultaneously characterize the sequence of personal genomes, resolving polymorphic SVs, together with their epigenomic status (Yue et al. 2022; Cheung et al. 2023; Sigurpalsdottir et al. 2024).

Moreover, computational methods that can compile personalized genomes into pangenome graphs, can capture megabases of non-reference sequences and integrate SVs from a cohort of genomes (Erik Garrison et al. 2023; Li, Feng, and Chu 2020; Hickey et al. 2023). The publication of the draft human pangenome reference also facilitates the study of SVs and their features at scale in a range of datasets (Liao et al. 2023; Groza et al. 2024). Indeed, such developments allow mapping epigenomic data directly to SVs and exploring the epigenetic status of regions that were not included in the reference genome (Groza et al. 2023; 2020). For example, it would be interesting to explore the link between SVs and 5mC base modifications, given the well known connection between DNA methylation and gene expression (Razin A and Cedar H 1991; Breiling and Lyko 2015; Dhar et al. 2021).

A pangenome can support genotyping SVs across the same samples or a wider set of samples, which fits well within the assumptions of most methylation QTL studies. Thus, mapping methylation data to pangenomes to correct reference bias and recover more signal (Wulfridge et al. 2019) and then correlating the resulting methylation features with SVs is a promising approach that is enabled by pangenomes. Therefore, the tools necessary to answer long-standing questions regarding the epigenetic status of SVs (Daron and Slotkin 2017; Groza et al. 2023; Sun et al. 2023) and their associations with other quantitative traits are increasingly accessible.

Here, we use a pangenome comprising 782 haplotype resolved de novo assemblies from the Genomic Answers for Kids (GA4K) Consortium (Cohen et al. 2022; Kane et al. 2023) and the 94 Human Pangenome Reference Consortium (HPRC) assemblies (Liao et al. 2023) to survey 435 5mC methylomes derived from whole genome sequencing of blood using HiFi long-reads. With this pangenomic approach, we identify non-reference CpGs within SVs, characterize their population frequency and methylation status, and associate SV-QTLs with methylation variation over the entire genome.

## Results

### Pangenomes characterize the methylation status of CpGs in SVs

We expected each of the 782 GA4K *de novo* assemblies to contain a number of structurally variant and non-reference CpGs. To recover these sequences, we constructed a genome graph using minigraph (Li, Feng, and Chu 2020) starting with CHM13v2 (Nurk Sergey et al. 2022) as a backbone, followed by the 94 HPRC assemblies (Liao et al. 2023), before adding the 782 GA4K samples. In total, this added 14.6 million CpGs in alternative nodes (SV-CpGs) for GA4K, on average 16,600 SV-CpGs per sample, on top of the 33.5 million CpGs that exist in CHM13v2 (a gain of 43.6%). At the same time, the pangenome grew by 713 megabases (Fig S1, a gain of 23%), yielding 2.05 × 10^4^ CpGs per megabase of non-reference sequence, compared to only 1.08 × 10^4^ CpGs per megabase of reference sequence.

Next, we obtained and aligned 435 GA4K blood methylomes matching a subset of the samples in this pangenome and annotated each CpG in the pangenome with the methylation level found in these samples (Methods). Overall, we found that these methylomes cover 7.99 million of the 14.6 million SV-CpGs that exist in the pangenome (Fig 1A). We also found that 5.6 million (64.8%) of these SV-CpGs show a methylation level above 50% (methylated state, SV-5mCpGs) in at least one of the 435 methylomes (Fig 1A). For comparison, 32.2 of 33.5 million reference CpGs (96.1%, Fig 1A) show a methylation level above 50% in the same samples (reference 5mCpGs). However, this difference was driven by rare SV-CpGs (Fig S2) that have few chances to be observed in a methylated state compared to reference CpGs that were nearly fixed in the population (Fig S3). In fact, 92.8% of the most common SV-CpGs were methylated in at least one sample (Fig S2). Saturation analysis over these methylomes shows that each additional methylome was expected to contribute 2860 SV-5mCpGs, 1750 SV-CpGs and 1700 reference 5mCpGs (Fig 1B). The fact that SV-CpGs saturated quicker than SV-5mCpGs also supports the hypothesis that more SV-CpGs should be observed in a methylated state as the number of methylomes increases.

**Fig 1:**
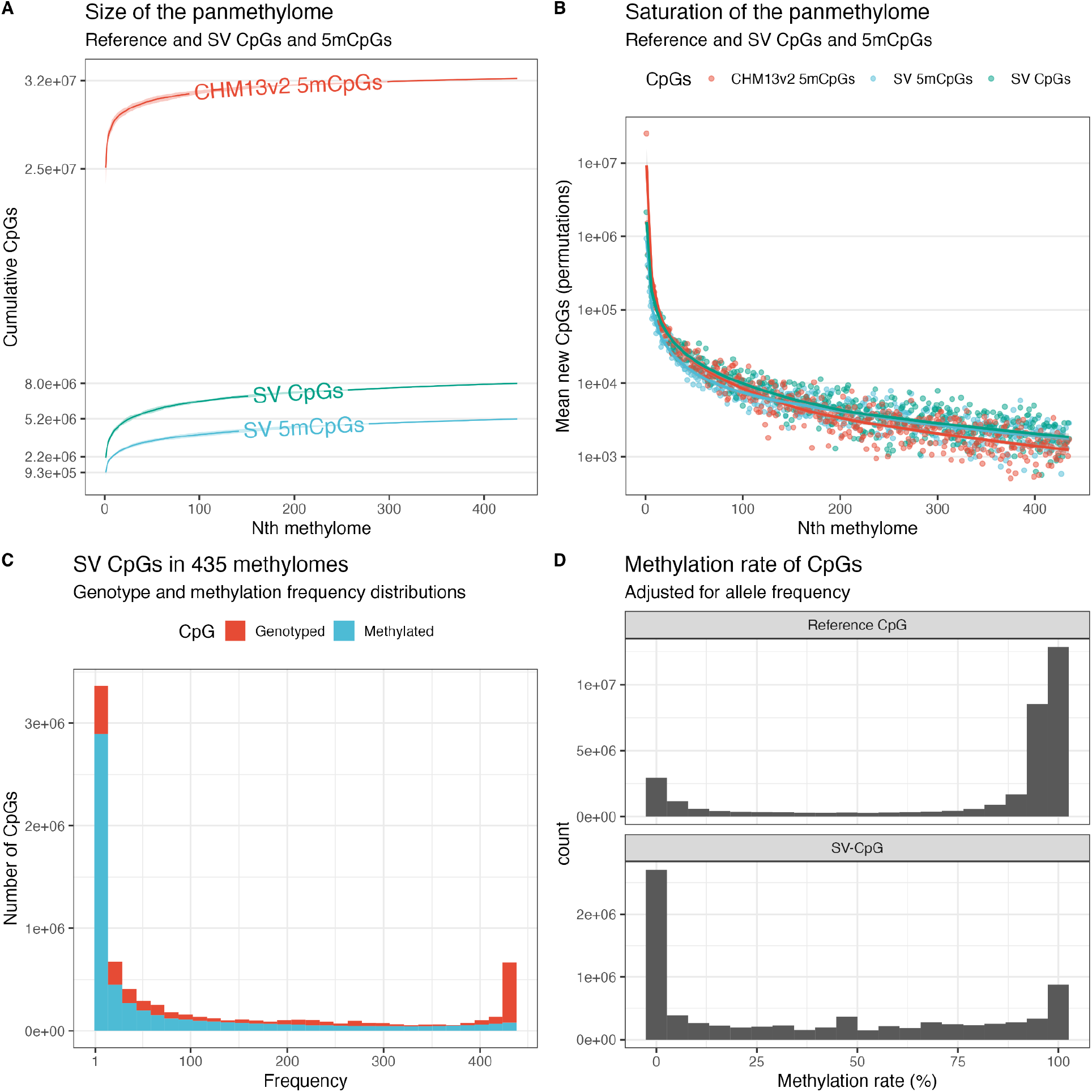
Number, frequency and methylation of non-reference CpGs. **A)** The total number of reference 5mCpGs, SV-CpGs, and SV-5mCpGs in the 435 methylomes. **B)** The rate of change in the saturation of CpGs shown in A). **C)** Frequency distribution of SV-CpGs (red) and SV-5mCpGs (blue). **D)** Observed methylation rates across SV- and reference CpGs, adjusted for allele frequency by counting only samples that carry any given CpG.

Using this approach, we were able to view methylation patterns in many haplotypes, including SVs, across hundreds of samples in polymorphic regions like the KIR (Fig 2). In this representation, we clearly see patterns of methylated and unmethylated CpGs within non-reference sequences in the KIR locus, a task that was not possible with reference genomes that describe only one haplotype.

**Fig 2:**
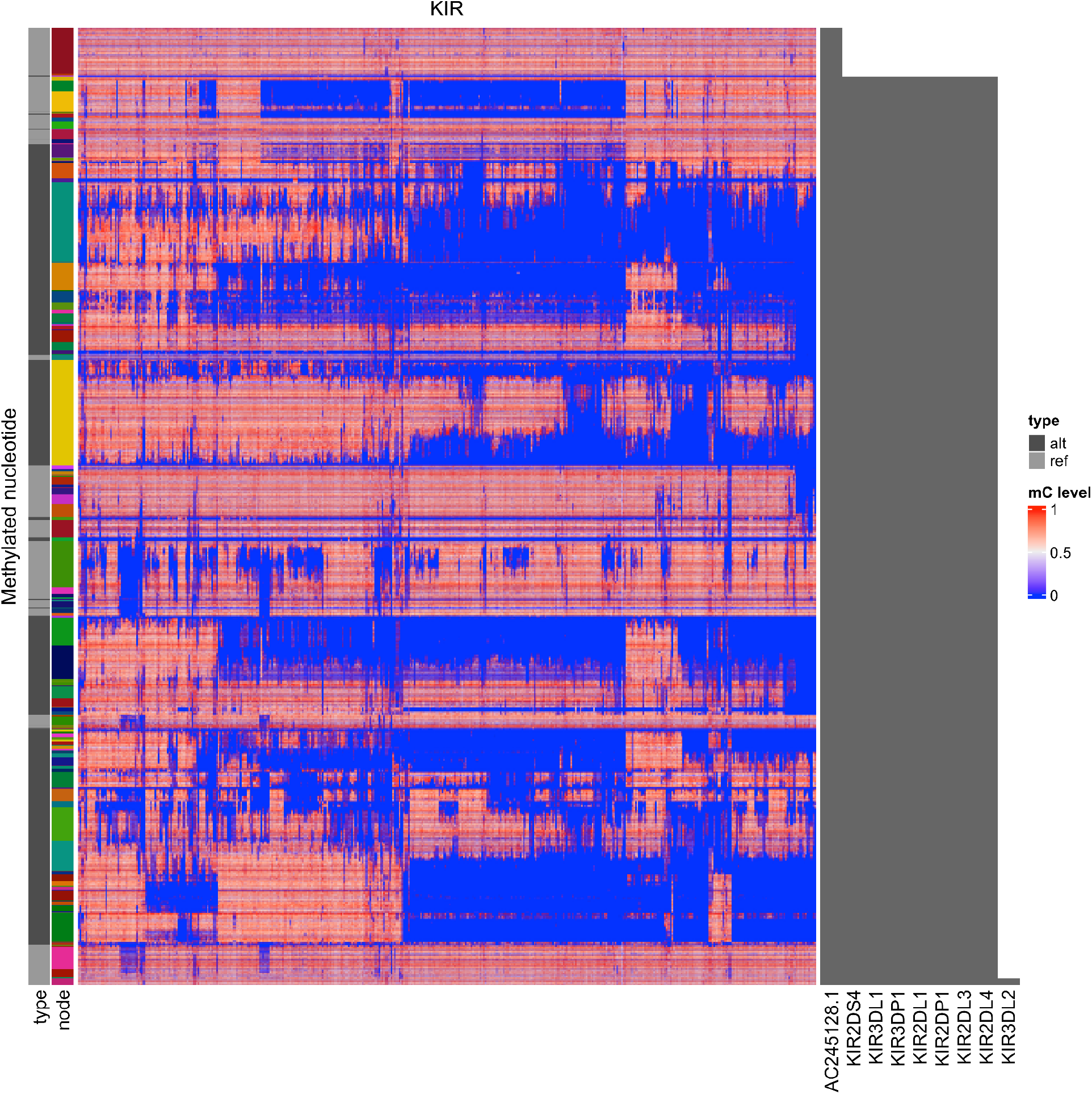
Heatmap visualization of methylation patterns in the KIR locus across 435 methylomes in the GA4K pangenome. Rows are CpGs and columns are methylomes. CpGs are ordered top to bottom, in the 5’ to 3’ direction as they appear in a haplotype, and are annotated by the node in the graph (the node row annotation) and whether it’s a reference or non-reference CpG (the type row annotation, alt or ref). The right annotation shows the genes that overlap the bubbles in which the CpGs lie.

### A large fraction of non-reference CpGs are methylated

Knowing the size of the above panmethylome, we asked what was the frequency distribution of SV-CpGs and SV-5mCpGs. We observe 8.48 × 10^5^ singleton SV-CpGs, 4.41 × 10^6^ with a frequency below 10% and 9.26 × 10^5^ with a frequency above 90% (Fig 1C). Some of these were methylated and we counted 9.48 × 10^5^ singleton SV-5mCpGs, 3.60 × 10^6^ with a frequency below 10% and 2.21 × 10^5^ with a frequency above 90% (Fig 1C). However, many 5mCpGs were rare because they lie on rare alleles. Therefore, we calculated the methylation rate, where we adjust for allele frequency and only consider samples that carry the CpG (Fig 1D, Methods). After we calculated methylation rates, we observed 2.4 × 10^6^ SV-CpGs (30.0%) and 7.1 × 10^5^ reference CpGs (2.14%) that were never methylated in any methylome and have a methylation rate of 0% (Fig 1D). Also, the average methylation rate was 38.9% for SV-CpGs and 76.6% for reference CpGs. Then, we counted CpGs that were fixed in a hypomethylated (<15%) or hypermethylated state (>85%) and found 3.44 × 10^6^ (43.1%) hypomethylated and 1.58 × 10^6^ (19.8%) hypermethylated SV-CpGs (Fig 1D). Conversely, we found 4.83 × 10^6^ (14.5%) hypomethylated and 2.34 × 10^7^ (70.4%) hypermethylated reference CpGs (Fig 1D). Lastly, we counted dynamic CpGs that have a methylation rate between 15% and 85%, yielding 2.97 × 10^6^ (37.2%) SV-CpGs and 4.93 × 10^6^ (14.8%) reference CpGs.

### Repeats account for more than half of the non-reference methylome

To determine the origin of SV-CpGs and SV-5mCpGs, we ran RepeatMasker on the pangenome and tallied the number of CpGs that overlap repeats. We found that 3.30 × 10^6^ SV-CpGs did not overlap any repeats (41.2%), of which 2.04 × 10^6^ (61.9%) were methylated at least once (Fig 3A). Second, centromeric satellites accounted for 1.58 × 10^6^ of SV-CpGs, of which 9.26 × 10^5^ (58.7%) were methylated. Mobile elements also contributed, with 6.67 × 10^5^ SV-CpGs in Alus (5.53 × 10^5^ methylated, 83.0%), 2.65 × 10^5^ in LINE1s (2.08 × 10^5^ methylated, 78.5%) and 1.87 × 10^5^ in SVAs (1.66 × 10^5^ methylated, 88.9%, Fig 3A).

**Fig 3:**
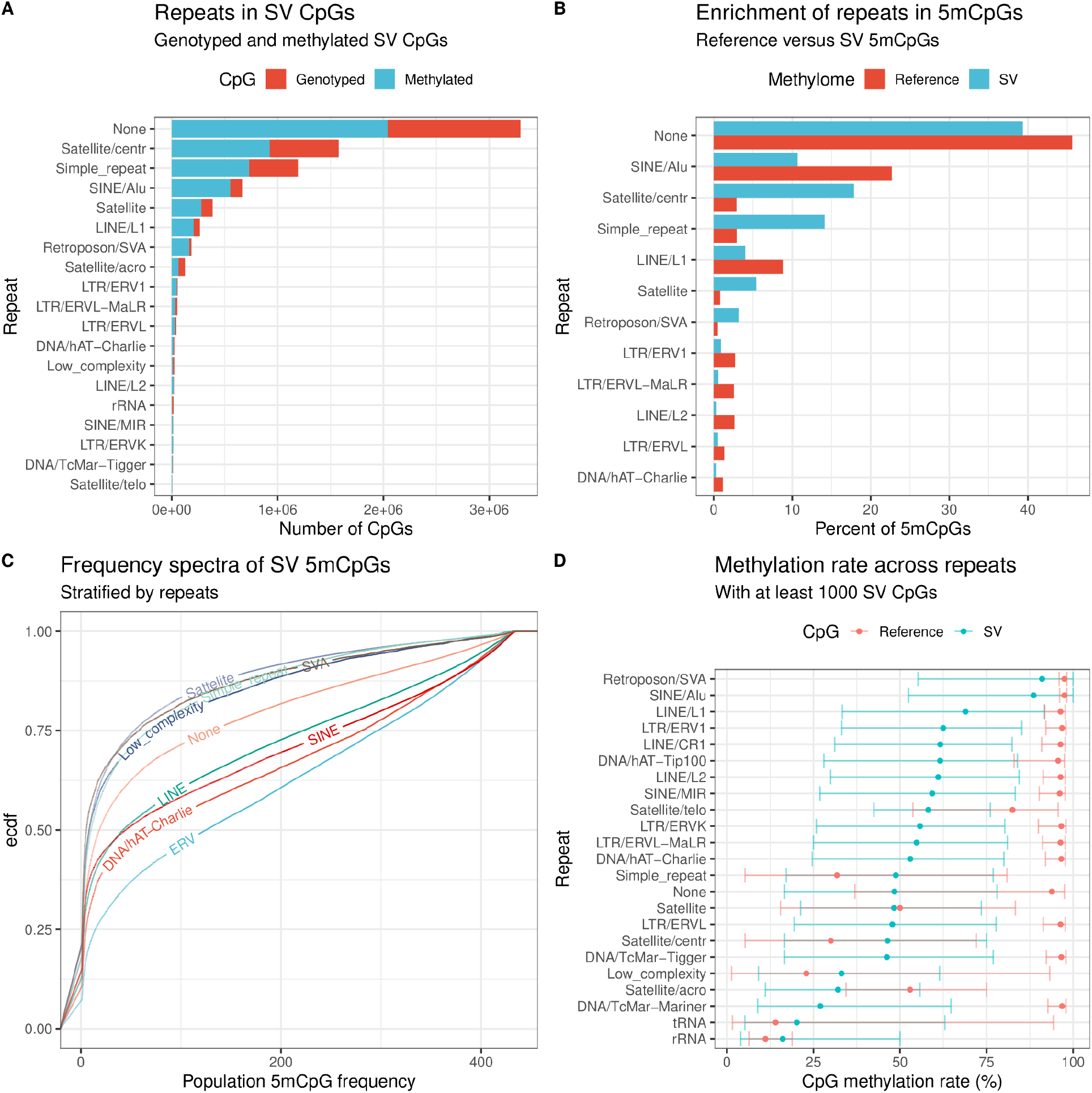
Annotation of sequences that contribute SV-CpGs. **A)** Number of SV-CpGs and SV-5mCpGs contributed by sequences without repeats (None) and sequences with repeats. **B)** Proportion of SV-5mCpGs contributed by each family of repeats, contrasted to the proportion of reference 5mCpGs contributed by the same repeats. **C)** Frequency distributions of SV-5mCpGs, stratified by repeat family. **D)** The observed methylation rates of 5mCpGs, as calculated in Fig 1D, stratified by repeat family. Intervals denote the 25%, 50% (median), and the 75% quantiles.

Then, we asked if any particular repeats contribute disproportionately to SV-5mCpGs relative to reference 5mCpGs. Here, we found that sequences that were not repeats were depleted in SV-5mCpGs, accounting for 45.6% of reference 5mCpGs but only for 39.4% of SV-5mCpGs. Similarly, Alus contribute 22.7% of reference 5mCpGs but only 10.7% of SV-5mCpGs. LINE1s, ERVs and other non-reference repeats were also depleted (Fig 3B). On the other hand, non-reference multi-allelic sequences such as satellites (0.835% vs 5.42%), centromeric satellites (2.91% vs 17.9%), simple repeats (2.94% vs 14.1%) and SVAs (0.514% vs 3.21%, known to contain tandem repeats) were overrepresented in SV-5mCpGs (Fig 3B).

Indeed, when we computed the frequency distribution of SV-5mCpGs and stratified by repeats (Fig 3C), we noted that SV-5mCpGs in overrepresented repeats like satellites, simple repeats and SVAs, were rarer than those in underrepresented repeats, which was consistent with multi-allelic SVs contributing a large number of rare alleles to the pangenome.

Finally, we wanted to know if the methylation rate of SV-CpGs (Fig 1D) varies across repeats and if it differs from reference CpGs. For this purpose, we calculated the methylation rate of every CpG in the pangenome and stratified by repeats, separating SV-CpGs from reference CpGs (Methods). In these analyses, we observed the methylation rate of SV-CpGs was highest in SVAs, Alus, LINE1s and ERV1s (Fig 3D). However, the methylation rate of SV-CpGs in these repeats tends to be lower and more variable than reference CpGs, pointing to a lower methylation rate in younger CpGs. Interestingly, the methylation rate of SV-CpGs in simple repeats, centromeric satellites, low complexity and tRNA sequences was higher than reference CpGs, and with similar variability (Fig 3D). In fact, these were multi-allelic regions where the reference genome often presents rare alleles which were also enriched in unmethylated CpGs (Fig S2). This suggests that the estimates of methylation rate were more variable in rarer CpGs due to low sample size. At the same time, the average methylation rate in rare SV-CpGs was lower than rare reference CpGs but is the same in common SV and reference CpGs (Fig S4). Given that these rare SV CpGs tend to lie in multi-allelic repeats, this bias could be explained by the lower mappability of these sequences. Moreover, there were very few rare reference CpGs (Fig S3), meaning that their methylation rate was estimated from a much smaller sample than for rare SV-CpGs.

### Pangenomes enable the mapping of SV-QTLs

We were interested to see if any SVs contained in the GA4K pangenome were QTLs for DNA methylation in our methylomes. To this end, we aligned and genotyped against the pangenome 470 haplotype-resolved *de novo* assemblies that were associated with the subset of 235 methylomes for which assemblies were available, selecting SV alleles that had a minimum frequency of 5% and a maximum frequency of 95% and resulted into a total of 160,064 SV alleles (Methods). The SV alleles were distributed in 97,746 loci, highlighting the ability of pangenomes to characterize multi-allelic regions. We also partitioned the CHM13v2 backbone reference into non-overlapping 200 bp methylation bins and calculated the average methylation level in these bins (Methods). Then, we identified SV alleles and methylation bins within 100kbp flanking sequence and performed 124.9 × 10^6^ SV-mQTL tests (Fig S5). We detected 230,464 methylation bins that were in QTL with 76,677 SV alleles in 59,872 loci at FDR < 0.05 (Fig 4A, example QTL in Fig 4B, Methods).

**Fig 4:**
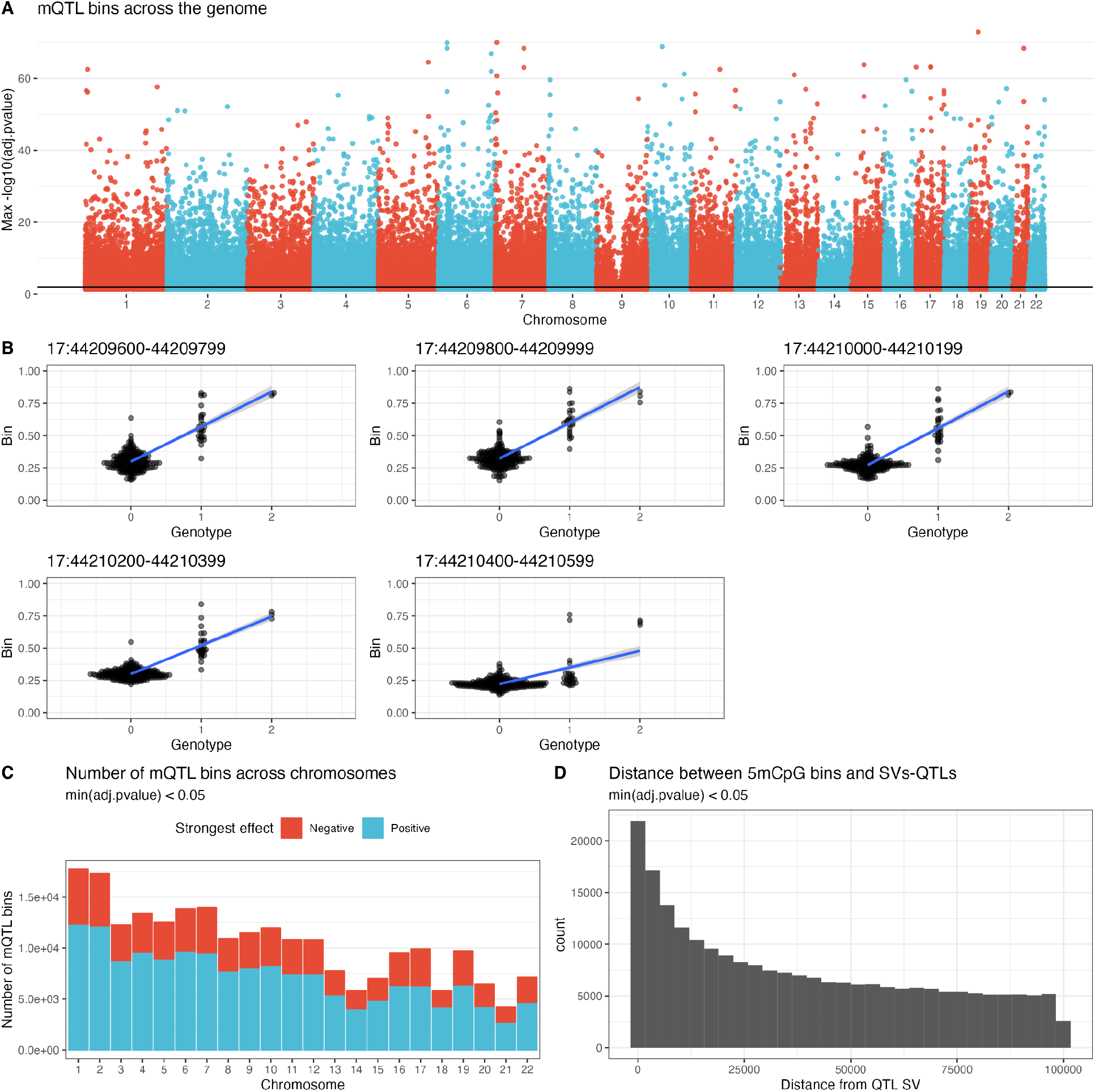
Quantitative trait locus mapping of 5mCpGs averaged in 200 bp methylation bins. **A)** Manhattan plot of QTL methylation bins associated with an SV at FDR < 0.05 over the entire reference genome backbone. **B)** Example of a leading SV-QTL interacting with the methylation of 5 consecutive bins. The nearest bin is 34 bp away from the SV. The SV allele is a 2216 bp deletion of CHM13v2.0#chr17:44210400-44210599. **C)** Number of methylation bins in QTL with an SV in the GA4K pangenome across chromosomes, stratified by positive and negative effect of the SV on methylation. **D)** Distribution of distances between SV-mQTLs and their methylation bins.

To describe the direction of mQTL effects for SVs, i.e. increase or decrease methylation, we tallied the signs of the strongest effect on each bin across chromosomes and found that SV-mQTLs showed more positive effects than negative effects: 73,702 bins were hypomethylated by SVs and and 156,762 were hypermethylated by the strongest SV-mQTL (Fig 4C). Every chromosome showed more hypomethylating than hypomethylating effect, with some chromosomes contributing slightly more QTLs relative to their size.

Next, we queried the distance distribution between the methylation bins and the associated SV-mQTLs to determine the ranges of interaction between SVs and methylation bins (Fig 4D). We tallied 4,144 methylation bins that overlap with their SV-mQTL, meaning the bin was within a structurally variant region of the backbone reference genome, and another 53,460 bins within 10 kbp of their SV-mQTL. Moreover, we found 14,970 methylation bins that were more than 90 kbp away from their SV-mQTL, at the limit of the allowed flanking distance. Overall, we observed a mean distance of interaction of 37.8 kbp (median 31.9 kbp).

We also wanted to know the methylation state of the SV-QTL alleles since it could be related to their QTL activity. Among the 55,149 SV alleles represented by paths containing at least one node in the pangenome graph, 22,125 alleles did not contain CpGs. In the remaining 33,024 SV alleles with CpGs, the average methylation rate was high, with 23,392 SVs having an average methylation rate above 85% (Fig S6). Moreover, reference and non-reference SV alleles have similar average methylation rate distributions.

### Some SVs are stronger methylation QTLs than SNPs

SVs are thought to be enriched in QTLs and have higher effect sizes (Jakubosky et al. 2020). To explore this hypothesis in the GA4K methylomes, we mapped SNP-mQTLs with the same parameters and frequency constraints as SV-mQTLs (Methods). In total, we tested 5,617,307 SNPs for associations with the same methylation bins and found 156,047 SNP-mQTL associated with 178,709 methylation bins at FDR < 0.05 (Table S1). Despite SNP-mQTLs being more numerous than SV-mQTLs, individual SVs were an order of magnitude more likely (17.2x) to be associated with methylation: only 2.78% of tested SNPs were mQTLs, in contrast to 47.9% of tested SVs. Then, we checked how often SVs were the leading variants over SNPs in mQTLs. For 65,659 methylation bins, the leading variant was an SV with a larger absolute effect size than any SNP (Fig 5A, Fig S7, example in Fig 4B). Conversely 145,453 SNPs were the top variant in 166,317 methylation bins. In terms of variants, 32,947 SV alleles out of 160,064 (20.6%) were the leading variant, compared to only 2.59% for SNPs, or a 7.95 fold enrichment (Table S1). Moreover, we found that SV-QTLs interact over longer distances (mean 37.7 kbp) than SNP-QTLs (mean 19.8 kbp) with methylation bins (Fig S8) and that each SV-QTL affects more methylation bins (mean 2.52 bins) than each SNP-QTL (mean 1.15 bins, Fig S9). For example, the SV-QTL in Fig 4B affects up to 5 methylation bins.

**Fig 5:**
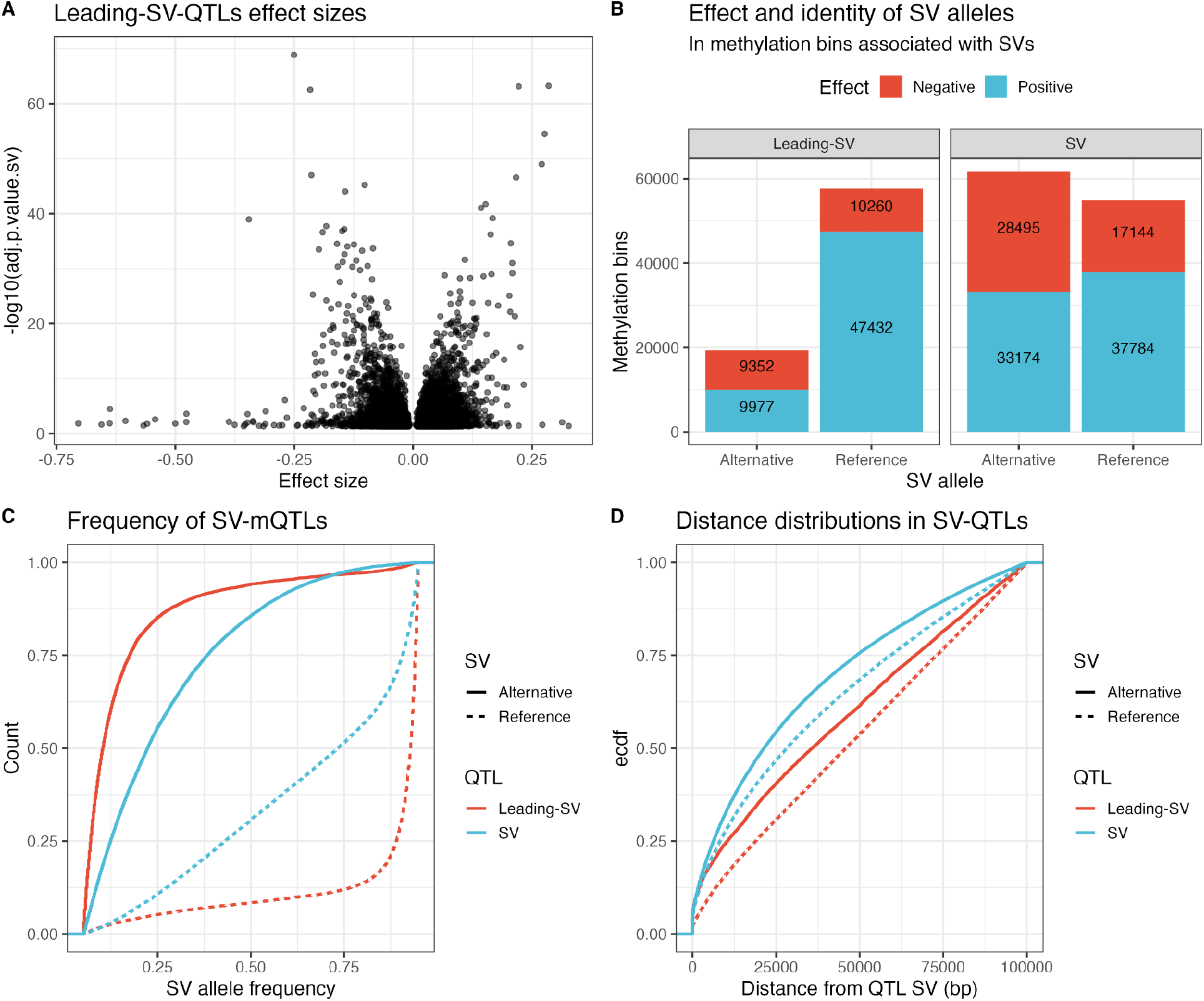
Leading mQTL variants among the pangenome SVs. **A)** Volcano plots of mQTLs where the leading variant is an SV. **B)** Number of methylation bins in mQTL with an SV, stratified by reference or non-reference SV allele, and positive or negative effect on methylation. **C)** Allele frequency distribution of mQTL-SVs, stratified by leading vs non-leading variant, and reference and non-reference SV alleles. **D)** The distribution of distances between SV-mQTLs and their methylation bins, stratified as in C).

### SV-QTLs are enriched in common SVs

When we looked at methylation bins where the leading variant was an SV, we noted that 57,692 bins were QTL with reference alleles having a predominantly positive effect on methylation (Fig 5B left). Another 19,329 bins were QTL with non-reference SV alleles, where positive and negative effects were evenly distributed. Next, we looked at methylation bins that were QTL with SVs but the leading variant was a SNP. Here, we found 61,669 methylation bins were QTL with non-reference SV alleles and 54,928 were QTL with reference SV alleles (Fig 5B right). Again, reference SV alleles show more positive effects on methylation. Thus, reference SV alleles tend to increase methylation and affect wider regions. More precisely, reference SV alleles were QTL with 2.14 bins on average, while non-reference SVs were QTL with only 1.70 methylation bins.

To explain these patterns, we plotted the frequency spectra of the SVs that were QTL with the above bins and found that leading reference SV alleles were very common, while leading non-reference SV alleles were the rarest (Fig 5C). To a lesser extent, the same pattern occurs with reference and non-reference SV alleles that were not leading variants (Fig 5C). We also explored if the ranges of interaction within QTLs were different between reference and non-reference SV alleles, and between SV alleles that were leading variants and those that were surpassed by SNPs. Here, we found that reference SV alleles were mQTL with methylation bins over longer distances than non-reference SV alleles (Fig 5D). Similarly, leading SV alleles were QTL over longer distances than SVs surpassed by SNPs. Lastly, repeat annotation of SV-QTL alleles again revealed that multi-allelic repeats were enriched in non-reference relative to reference SV alleles (Fig S10).

These differences in allele frequency and range of interaction suggest that the more frequent SV alleles interact with methylation more often and over longer distances than younger and less frequent SVs. At the same time, some rare SV alleles showed effects on methylation that were stronger than any nearby SNP.

### Leading SV-mQTLs are in proximity to GWAS SNPs

To highlight SVs that may be causal for methylation, we first identified SNVs that were in high linkage disequilibrium (R2 > 0.95) with previously known GWAS SNPs in the NHGRI-EBI GWAS Catalog (Sollis et al. 2023). Then, we filtered for SVs showing higher effect sizes than SNVs, that were closer to the methylation bin than the leading SNV and also no further than 10 kbp. In doing so, we obtained a list of 606 SVs than were QTL with 2,417 methylation bins (Table S2). That is, each putatively causal SV was in QTL on average with 3.99 methylation bins, which was 58% more than the average of SV-QTLs. Moreover, we annotated these SVs with 671 genes that were associated with GWAS SNPs in LD with GA4K SNVs (Table S2).

## Discussion

On average, we observed 16,600 new CpGs in the DNA sequences added by each genome to the GA4K pangenome (0.81 Mbp per genome). Moreover, we were able to characterize the repeat families and other sequences that contribute to new CpGs and showed that at least 64.8% of these new CpGs were methylated in at least one individual. Aided by the GA4K pangenome graph, we arranged and sorted this non-reference epigenetic variation in haplotypes that could be compared across many methylomes, allowing for characterization of complex patterns of methylations within SVs. Moreover, saturation analysis suggests that expanding this panmethylome with more assemblies and methylomes would continue to add thousands of new SV-CpGs and SV-5mCpGs per sample.

A minority of CpGs are known to be variable, or dynamic, in tissues (Ziller et al. 2013). Similarly we found that most CpGs in personal reference DNA show either predominant hypo -or hypermethylation and only 15% were variably methylated (15-85% methylated) in population haploid assemblies. In contrast, nearly 40% of non-reference DNA found in alternative nodes in the pangenome graph were variably methylated, showing they contribute disproportionately to epigenetic variation in humans. We could also use the pangenome graph to explore population variation in functional DNA. We could link nearly 60,000 SV loci with methylation variation in over 46.1 Mbp of DNA across population assemblies. Parallel analyses of SNV and SV variation in the same samples demonstrated larger prevalence of methylation among fewer SVs, their impacts extending greater distances and ability to explain a substantial proportion of mQTLs. We note that these comparisons did not include methylation bins that lie in new non-reference sequences not mappable by standard mQTL-SNV associations. Next, we identified two sets of leading SV-QTLs that surpass SNVs in effect size. First, we found a set of leading SV alleles that are common in the population. These tend to be positively associated with DNA methylation and are often included in the reference genome. Second, we found another set of rarer SV alleles that are associated equally with positive and negative effects on DNA methylation.

In conclusion, our observations underscore the importance of assessing genomes for the entirety of sequence space not only for structural but also for functional variation (Groza et al. 2023). We also confirm the properties of SVs linking to QTLs impacting at greater distances and at a higher frequency. Hypermethylation in regulatory elements canonically leads to loss of activity, which we previously exploited in rare variant characterization of long-read sequences (Cheung et al. 2023). The hypermethylating impact we observed for leading SV-mQTLs suggests an important role for studying SVs in gene silencing. Overall, the full scope structural variation catalogued in pangenome graphs suggests large utility in quantitative trait and disease genetic studies.

## Methods

### Creating the pangenome graph

We created the GA4K pangenome using `minigraph -xcggs –ggen` (version 0.20-r559). We started with the CHM13v2 backbone reference, added 94 HPRC genomes and finally 782 GA4K assemblies.

### Genotyping assemblies

To genotype SVs in the GA4K assemblies, we aligned them to the final pangenome and called variants using `minigraph -c –call –vc.` Then we created a unified genotype matrix across genomes by considering each bubble start, end, source, sink and path in the pangenome as alleles and then listing the presence of an allele in a genome as 1 and their presence as 0 (see `genotypes.R`). Reference alleles were determined by genotyping CHM13v2 against the pangenome graph.

### Annotating CpGs in the pangenome with repeats

We ran RepeatMasker with the Dfam_2.0 (Hubley et al. 2016) database on all nodes in the graph to identify repeats. Then, we overlapped the repeat annotation of nodes with the position of CpGs on these nodes to label each CpG with a repeat annotation.

### Mapping methylation data to pangenome graphs

We wrote `panmethyl` (https://github.com/cgroza/panmethyl) to map methylation data to pangenome graphs. To do so, `panmethyl` takes BAMs processed and annotated with MM and ML tags by PacBio Jasmine as input to extract HiFi long-reads that are annotated with methylation probabilities at each cytosine. Then, the reads are aligned to the pangenome with `minigraph --vc -c -N 1` and the methylation probabilities of every cytosine in the read is lifted to their corresponding positions in the pangenome graph. To calculate the per sample methylation level of cytosines in the pangenome, we averaged the methylation probabilities that map to their respective position. We also indexed every CpG in the pangenome and named CpG that are not in CHM13v2 as SV-CpGs. Using this index, we calculated the population frequency of reference CpGs and SV-CpGs by counting the number of methylomes that cover the CpG in the aligned reads. To calculate the population frequency of reference 5mCpGs and SV-5mCpGs, we counted the number of methylomes where the CpG has an average methylation level across strands of 50% or more (see `merge_cpgs.py`). We intersected CpGs with repeats to obtain the population frequency of CpGs and 5mCpGs stratified by repeats.

### Calculating the size and saturation of the panmethylome

We genotyped the presence and absence of CpGs by listing the nodes covered by the aligned methylomes. To obtain saturation curves for the panmethylome, we simulated 10 curves, each with a randomly permuted order of methylomes. In each permutation, we start with the first methylome and progressively add newly discovered SV-CpGs, SV-5mCpGs and 5mCpGs in subsequent methylomes (see `cpg_saturation.py`). At every iteration, we record the number of accumulated CpGs and the number of new CpGs. To extrapolate the saturation rate, we fit a logarithmic function on the number of new CpGs across the average of the 10 simulations.

### Calculating the methylation rate of CpGs

To know how often each CpG is methylated, we calculated the methylation rate as follows:

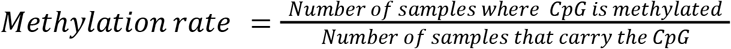

We intersected the CpGs with repeat annotations, to obtain methylation rates stratified by repeats.

### Mapping methylation QTLs

To map methylation SV-mQTLs and SNP-mQTLs, we binned the CHM13v2 backbone reference genome into non-overlapping bins that are 200 bp in length. Then we averaged the methylation levels of every cytosine in each bin. Cytosines with missing data are not considered in the average methylation level (see `bin_methylation.R`). To map QTLs for each methylation bin, we ran linear regression using `lm()` between the methylation level of the bin and the genotype of every SV within 100 kbp (see `run_qtls.R)`. Here, the genotype was the number of SV alleles (0, 1, 2) carried by each sample. We did the same for methylation bins and every SNP within 100 kbp. We corrected for multiple testing using `p.adjust(method = “fdr”)`. We selected the SV with the lowest significant (0.05 FDR) adjusted p-value ber bin. To rank and find the leading SV-mQTLs and SNP-mQTLs, we compared the absolute effect sizes of every variant associated with a methylation bin with an FDR < 0.05.

## Acknowledgements

We would like to thank all families for participating in the Genomic Answers for Kids study. This work was made possible by generous gifts to the Children’s Mercy Research Institute and Genomic Answers for Kids program at Children’s Mercy Kansas City. We also would like to thank Nick Nolte, Dan Louiselle, and Rebecca Biswell for their work in sample processing, Laura Puckett and Adam Walters for their work in library preparation and sequencing, and the clinical coordination team led by Bradley Belden for their work in clinical coordination. We also would like to thank PacBio for sequencing support for a subset of the samples. T.P. holds the Dee Lyons/Missouri Endowed Chair in Pediatric Genomic Medicine, and E.G., holds the Roberta D. Harding & William F. Bradley, Jr. Endowed Chair in Genomic Research. C.G. is supported by the NSERC PGS D award. G.B. is supported by a Canada Research Chair Tier 1 award and an FRQ-S Distinguished Research Scholar award. This research was enabled in part by support provided by Calcul Quebec and the Digital Research Alliance of Canada.

## Data availability

The 5-base HiFi-GS, HiFi long-read transcript sequencing (IsoSeq) and WGBS raw and processed data including assemblies and genotypes generated in this study have been deposited in the dbGAP (https://www.ncbi.nlm.nih.gov/gap/) database under accession code phs002206.v4.p1 [https://www.ncbi.nlm.nih.gov/projects/gap/cgi-bin/study.cgi?study_id=phs002206.v4.p1]. Raw and processed data are available under restricted access due to IRB regulations and informed consent limiting access to users studying genetic diseases. Data access is provided by dbGAP (https://dbgap.ncbi.nlm.nih.gov/aa/wga.cgi?page=login) for certified investigators with local IRB approval in place. The CHM13v2.0 reference genome is available for download at https://s3-us-west-2.amazonaws.com/human-pangenomics/T2T/CHM13/assemblies/analysis_set/chm13v2.0.fa.gz. Methylation counts are available at https://zenodo.org/doi/10.5281/zenodo.10825920.

## Code availability

Panmethyl is available at https://github.com/cgroza/panmethyl. Scripts are available at https://zenodo.org/doi/10.5281/zenodo.10825920.

## Notes

### Competing Interest Statement

The authors have declared no competing interest.

## References

Breiling, Achim, and Frank Lyko. 2015. “Epigenetic Regulatory Functions of DNA Modifications: 5-Methylcytosine and Beyond.” Epigenetics & Chromatin 8 (1): 24. 10.1186/s13072-015-0016-6.

Cheung, Warren A., Adam F. Johnson, William J. Rowell, Emily Farrow, Richard Hall, Ana S. A. Cohen, John C. Means, et al. 2023. “Direct Haplotype-Resolved 5-Base HiFi Sequencing for Genome-Wide Profiling of Hypermethylation Outliers in a Rare Disease Cohort.” Nature Communications 14 (1): 3090. 10.1038/s41467-023-38782-1.

Cohen, Ana S. A., Emily G. Farrow, Ahmed T. Abdelmoity, Joseph T. Alaimo, Shivarajan M. Amudhavalli, John T. Anderson, Lalit Bansal, et al. 2022. “Genomic Answers for Children: Dynamic Analyses of >1000 Pediatric Rare Disease Genomes.” Genetics in Medicine : Official Journal of the American College of Medical Genetics 24 (6): 1336–48. 10.1016/j.gim.2022.02.007.

Daron, Josquin, and R. Keith Slotkin. 2017. “EpiTEome: Simultaneous Detection of Transposable Element Insertion Sites and Their DNA Methylation Levels.” Genome Biology 18 (1): 91. 10.1186/s13059-017-1232-0.

Dhar, Gaurab Aditya, Shagnik Saha, Parama Mitra, and Ronita Nag Chaudhuri. 2021. “DNA Methylation and Regulation of Gene Expression: Guardian of Our Health.” The Nucleus 64 (3): 259–70. 10.1007/s13237-021-00367-y.

Erik Garrison, Andrea Guarracino, Simon Heumos, Flavia Villani, Zhigui Bao, Lorenzo Tattini, Jörg Hagmann, et al. 2023. “Building Pangenome Graphs.” bioRxiv, January, 2023.04.05.535718. 10.1101/2023.04.05.535718.

Gershman, Ariel, Michael E. G. Sauria, Xavi Guitart, Mitchell R. Vollger, Paul W. Hook, Savannah J. Hoyt, Miten Jain, et al. 2022. “Epigenetic Patterns in a Complete Human Genome.” Science 376 (6588): eabj5089. 10.1126/science.abj5089.

Groza, Cristian, Xun Chen, Alain Pacis, Marie-Michelle Simon, Albena Pramatarova, Katherine A. Aracena, Tomi Pastinen, Luis B. Barreiro, and Guillaume Bourque. 2023. “Genome Graphs Detect Human Polymorphisms in Active Epigenomic State during Influenza Infection.” Cell Genomics, May. 10.1016/j.xgen.2023.100294.

Groza, Cristian, Tony Kwan, Nicole Soranzo, Tomi Pastinen, and Guillaume Bourque. 2020. “Personalized and Graph Genomes Reveal Missing Signal in Epigenomic Data.” Genome Biology 21 (1): 124. 10.1186/s13059-020-02038-8.

Groza, Cristian, Carl Schwendinger-Schreck, Warren A. Cheung, Emily G. Farrow, Isabelle Thiffault, Juniper Lake, William B. Rizzo, et al. 2024. “Pangenome Graphs Improve the Analysis of Structural Variants in Rare Genetic Diseases.” Nature Communications 15 (1): 657. 10.1038/s41467-024-44980-2.

Hickey, Glenn, Jean Monlong, Jana Ebler, Adam M. Novak, Jordan M. Eizenga, Yan Gao, Haley J. Abel, et al. 2023. “Pangenome Graph Construction from Genome Alignments with Minigraph-Cactus.” Nature Biotechnology, May. 10.1038/s41587-023-01793-w.

Hubley, Robert, Robert D Finn, Jody Clements, Sean R Eddy, Thomas A Jones, Weidong Bao, Arian F A Smit, and Travis J Wheeler. 2016. “The Dfam Database of Repetitive DNA Families.” Nucleic Acids Research 44 (D1): D81–89. 10.1093/nar/gkv1272.

Jakubosky, David, Matteo D’Antonio, Marc Jan Bonder, Craig Smail, Margaret K. R. Donovan, William W. Young Greenwald, Hiroko Matsui, et al. 2020. “Properties of Structural Variants and Short Tandem Repeats Associated with Gene Expression and Complex Traits.” Nature Communications 11 (1): 2927. 10.1038/s41467-020-16482-4.

Kane, N. J., A. S. A. Cohen, C. Berrios, B. Jones, T. Pastinen, and M. A. Hoffman. 2023. “Committing to Genomic Answers for All Kids: Evaluating Inequity in Genomic Research Enrollment.” Genetics in Medicine : Official Journal of the American College of Medical Genetics, May, 100895. 10.1016/j.gim.2023.100895.

Li, Heng, Xiaowen Feng, and Chong Chu. 2020. “The Design and Construction of Reference Pangenome Graphs with Minigraph.” Genome Biology 21 (1): 265. 10.1186/s13059-020-02168-z.

Liao, Wen-Wei, Mobin Asri, Jana Ebler, Daniel Doerr, Marina Haukness, Glenn Hickey, Shuangjia Lu, et al. 2023. “A Draft Human Pangenome Reference.” Nature 617 (7960): 312–24. 10.1038/s41586-023-05896-x.

Nurk Sergey, Koren Sergey, Rhie Arang, Rautiainen Mikko, Bzikadze Andrey V., Mikheenko Alla, Vollger Mitchell R., et al. 2022. “The Complete Sequence of a Human Genome.” Science 376 (6588): 44–53. 10.1126/science.abj6987.

Razin A and Cedar H. 1991. “DNA Methylation and Gene Expression.” Microbiological Reviews 55 (3): 451–58. 10.1128/mr.55.3.451-458.1991.

Sigurpalsdottir, Brynja D., Olafur A. Stefansson, Guillaume Holley, Doruk Beyter, Florian Zink, Marteinn Þ. Hardarson, Sverrir Þ. Sverrisson, et al. 2024. “A Comparison of Methods for Detecting DNA Methylation from Long-Read Sequencing of Human Genomes.” Genome Biology 25 (1): 69. 10.1186/s13059-024-03207-9.

Sollis, Elliot, Abayomi Mosaku, Ala Abid, Annalisa Buniello, Maria Cerezo, Laurent Gil, Tudor Groza, et al. 2023. “The NHGRI-EBI GWAS Catalog: Knowledgebase and Deposition Resource.” Nucleic Acids Research 51 (D1): D977–85. 10.1093/nar/gkac1010.

Sun, Zhifu, Saurabh Behati, Panwen Wang, Aditya Bhagwate, Samantha McDonough, Vivian Wang, William Taylor, Julie Cunningham, and John Kisiel. 2023. “Performance Comparisons of Methylation and Structural Variants from Low-Input Whole-Genome Methylation Sequencing.” Epigenomics 15 (1): 11–19. 10.2217/epi-2022-0453.

Wulfridge, Phillip, Ben Langmead, Andrew P Feinberg, and Kasper D Hansen. 2019. “Analyzing Whole Genome Bisulfite Sequencing Data from Highly Divergent Genotypes.” Nucleic Acids Research 47 (19): e117–e117. 10.1093/nar/gkz674.

Yue, Xue, Zhiyuan Xie, Moran Li, Kai Wang, Xiaojing Li, Xiaoqing Zhang, Jian Yan, and Yimeng Yin. 2022. “Simultaneous Profiling of Histone Modifications and DNA Methylation via Nanopore Sequencing.” Nature Communications 13 (1): 7939. 10.1038/s41467-022-35650-2.

Ziller, Michael J., Hongcang Gu, Fabian Müller, Julie Donaghey, Linus T.-Y. Tsai, Oliver Kohlbacher, Philip L. De Jager, et al. 2013. “Charting a Dynamic DNA Methylation Landscape of the Human Genome.” Nature 500 (7463): 477–81. 10.1038/nature12433.

